# Monotonous behavior with 2-propanol converts into reentrant transition with 1-propanol: Higher-order structure of DNA

**DOI:** 10.1101/715391

**Authors:** Y. Ma, Y. Yoshikawa, K. Yoshikawa

## Abstract

In the present study, we measured the changes in the higher-order structure of genomic DNA molecules in the presence of alcohols through single-DNA observation by use of fluorescence microscopy, with particular focus on the different effects of 1-propanol and 2-propanol. The results showed that, with an increasing concentration of 1-propanol, DNA exhibits reentrant conformational transitions from an elongated coil to a folded globule, and then to an unfolded state. On the other hand, with 2-propanol, DNA exhibits monotonous shrinkage into a compact state. Thus, DNA molecules are more effectively condensed/precipitated with 2-propanol than with 1-propanol. The propanol isomers also had different effects on the changes in the secondary structure of DNA, as revealed by circular dichroism (CD) measurements. With 1-propanol, DNA maintains a B-form secondary structure. An A-like conformation appears with the addition of 2-propanol.

**STATEMENT OF SIGNIFICANCE:** Currently, 2-propanol has most often been used as the solvent to extract and purify genomic DNA molecules from living cells, according the protocols in molecular biology and biochemistry. Unfortunately, the reason why usage of 2-propanol is recommended instead of ethanol and 1-propanol has never been explained in a clear manner. We believe that the new insight based on chemical physics point of view would play an important role for the development of current chemical procedures/treatments adapted on an empirical basis.

## INTRODUCTION

In biomedical experiments, alcohol precipitation is widely used to purify or concentrate nucleic acids.(1–11) 2-Propanol (isopropyl alcohol) is often used to isolate DNA molecules from cells through precipitation.(4, 6),(7, 9, 10) On the other hand, 1-propanol is not adapted as the solvent for DNA precipitation. Currently, 1-propanol is regarded as a useful solvent in the pharmaceutical industry, and is used mainly for resins and cellulose esters.(12, 13) Although 2- and 1-propanol are used for different purposes, the underlying physicochemical difference between these isomers seems to have not yet been clarified.

The structural transition of DNA between elongated coil and compacted globule states, the so-called coil-globule transition, is one of the central problems in the fields of biochemistry, biophysics, and soft matter physics. Several studies have examined this problem from both theoretical and experimental approaches.(14–26) Over the past couple of decades, it has been established that large DNA above the size of several tens of kilo base pairs (kbp) exhibits unique conformational characteristics, including the occurrence of discrete transition between elongated coil and compact state on individual DNA molecular chain, and also formation of intrachain segregation, as demonstrated by singlemolecule observation in bulk solutions using fluorescence microscopy. (19, 23, 26, 27)

The secondary structure of DNA is closely connected to its function. It has been reported that the secondary structure of DNA is a common causative factor in human disease that also influences human telomerase activity.(28, 29) Many studies have examined the secondary structure of DNA, such as A-, B-, C- and Z-forms, in different chemicals solutions, including alcohols.(30–35)In this study, the higher-order structure of giant DNA was evaluated through single-molecule observation using fluorescence microscopy.

## EXPERIMENTAL METHODOLOGY

### Materials and preparation of samples

The bacteriophage λ-DNA (48 kbp) was purchased from Nippon Gene. DNA samples were dissolved in propanol-water solution with a final concentration of 30 μM in nucleotide units. A fluorescent cyanine dye, YOYO-1 (quinolinium, 1, 1’-[1, 3-propanediyl-bis [(dimethylimino)-3, 1-propanediyl]] bis [4 - [(3 – methyl - 2(3H) - benzoxazolylidene) - methyl]] - tetraiodide), was purchased from Molecular Probes, Inc. (Eugene, OR, USA). The antioxidant 2-mercaptoethanol (2-ME), 1- and 2-propanol were purchased from Wako Pure Chemical Industries (Osaka, Japan). YOYO-1 (final concentration: 1μM) was added to the DNA solution, together with 4% (v/v) 2-ME.

### Observation of the higher-order structure of DNA by fluorescence microscopy

Fluorescence DNA images were captured using an Axiovert 135 TV microscope (Carl Zeiss, Oberkochen, Germany) equipped with an oil-immersed 100x objective lens, and recorded on DVD through an EBCCD camera (Hamamatsu Photonics, Hamamatsu, Japan). The recorded videos were analyzed using VirtualDub (written by Avery Lee) and ImageJ software (National Institute of Mental Health, Bethesda, MD, USA). All observations were carried out at around 25 °C.

### CD measurements

The CD spectra of λ-DNA were measured with a CD spectrometer (J-820, JASCO, Japan). Measurements were performed at a scan rate of 100 nm/min and 2000 μL of each sample was tested at around 25 °C. The cell path length was 1 cm and CD spectra were obtained as the accumulation of three scans.

## RESULTS

### Higher-order structure of DNA molecules in alcohol solutions

To evaluate the change of the higher-order structure of λ-DNA molecules in solution, we measured the long-axis length, L,(19, 23, 25, 26, 36, 37) as shown in the fluorescence images and schematics in Fig. 1. The fluorescence images [propanol concentrations from top to bottom: 0%, 50%, 70% (v/v) (1-propanol)] in Fig. 1 show three representative conformations of DNA molecules. Partial globule (19, 23, 26, 27) is the state where compact and elongated parts coexist along a single λ-DNA molecular chain.

**FIG. 1.**
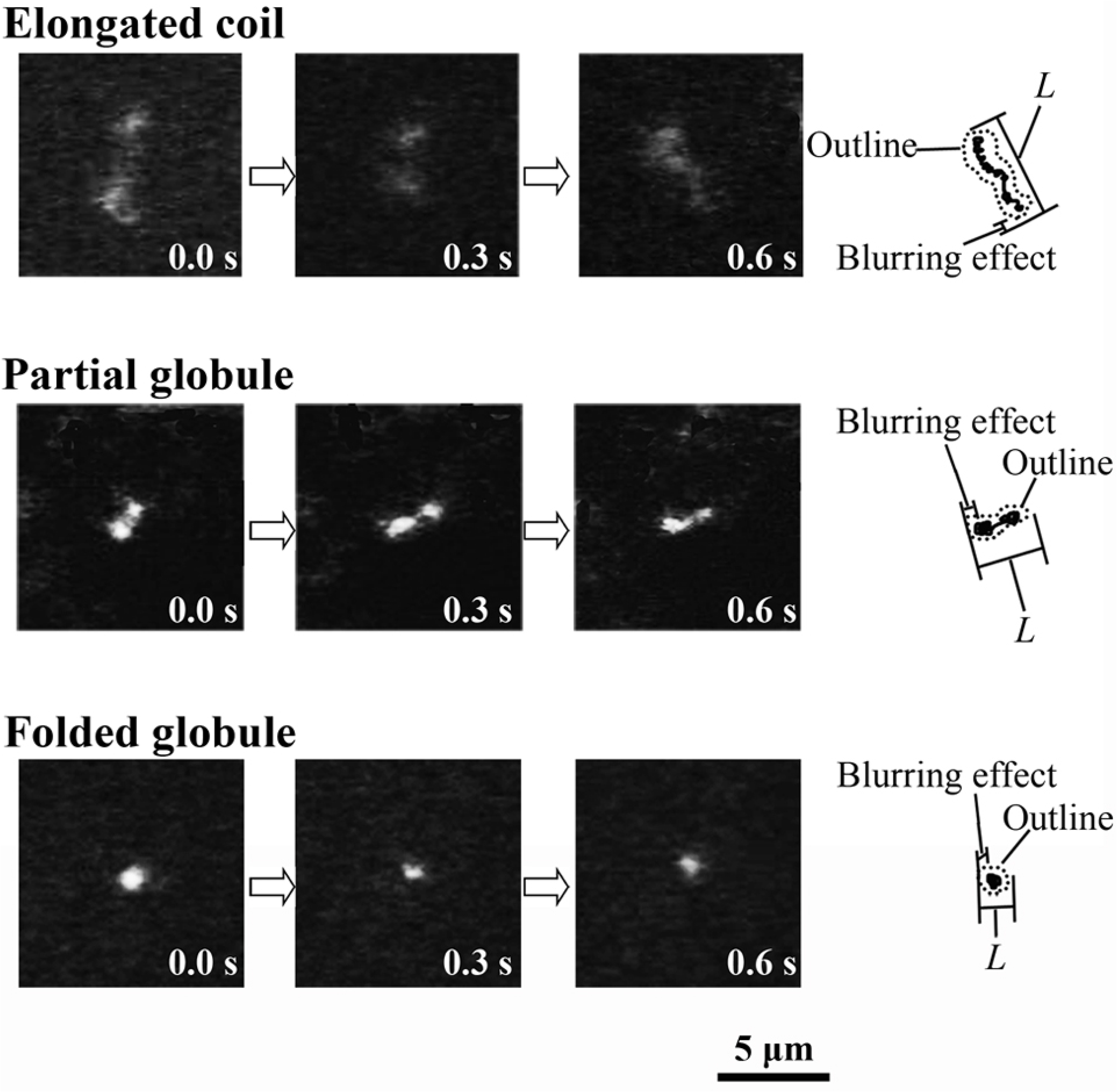
Fluorescence images of λ-DNA molecules (48 kbp) exhibiting Brownian motion in 1-propanol solutions; from top to bottom: 0, 50, 80 (v/v) %.

As shown in Fig. 2 a, after the DNA samples in 50(v/v)% 1-propanol are subjected to an electric field of about 10 V/cm,(23) the twin bright spots tend to be separated accompanied by the elongation of the interconnected coil part, revealing that folded compact and elongated coil parts coexist along a single DNA molecule, i.e., the existence of partial globule state.(23, 27) In contrast, in Fig. 2 b for the solution of 60(v/v)% 1-propanol, the bright spot does not separate after the application of an electric field, which demonstrates that the molecule is in a folded globule state.

**FIG. 2.**
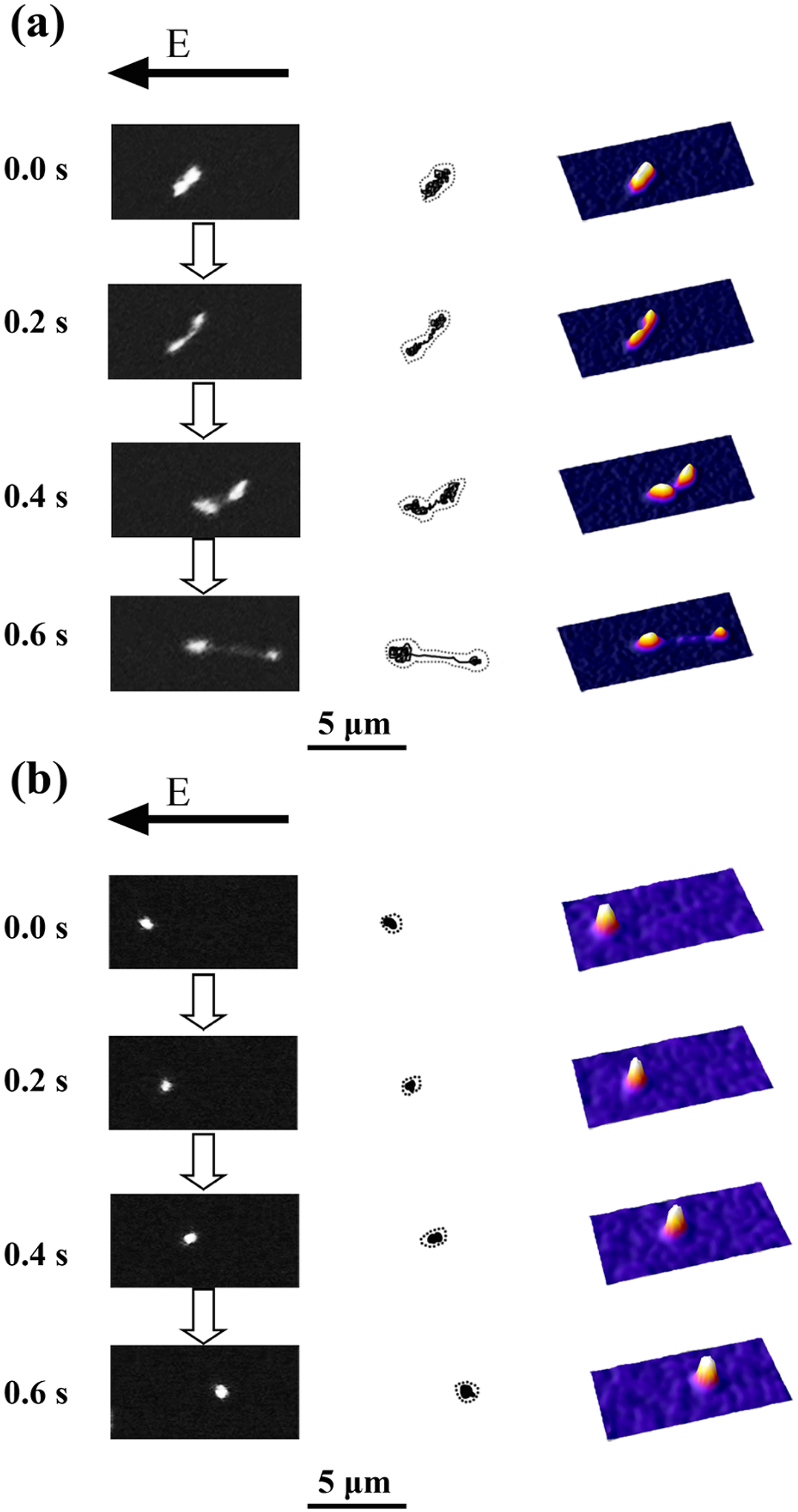
Change in the conformation of a single λ-DNA molecule after the application of a DC electric field, E, of ca.10 V/cm. Left; fluorescence images, Middle; schematics of DNA conformation, Right; quasi-three-dimensional representations on the fluoresce intensity distribution. (a) The partial globular molecule in 50(v/v)% 1-propanol solution; there is clear segregation of the globular and coil parts; (b) The folded globular molecule in 60(v/v)% 1-propanol solution.

Figure 3 shows the distribution of the long-axis length of DNA molecules. The long-axis length decreases with an increase in the concentration of propanol solutions. The average long-axis lengths of the correspond data are shown in Fig. 5 a. For DNA molecules in different concentrations of 1-propanol, one minimum appeared at 60 % (v/v), suggesting the occurrence of reentrant conformation transition on coil-globule-coil states. In 2-propanol solutions, the average long-axis length of DNA decreased, and a transition from an elongated coil state to a folded globule occurred at around 70-80% (v/v). Above 70% (v/v), DNA maintained a folded globule conformation and the average length of DNA molecules remained essentially constant.

**FIG. 3.**
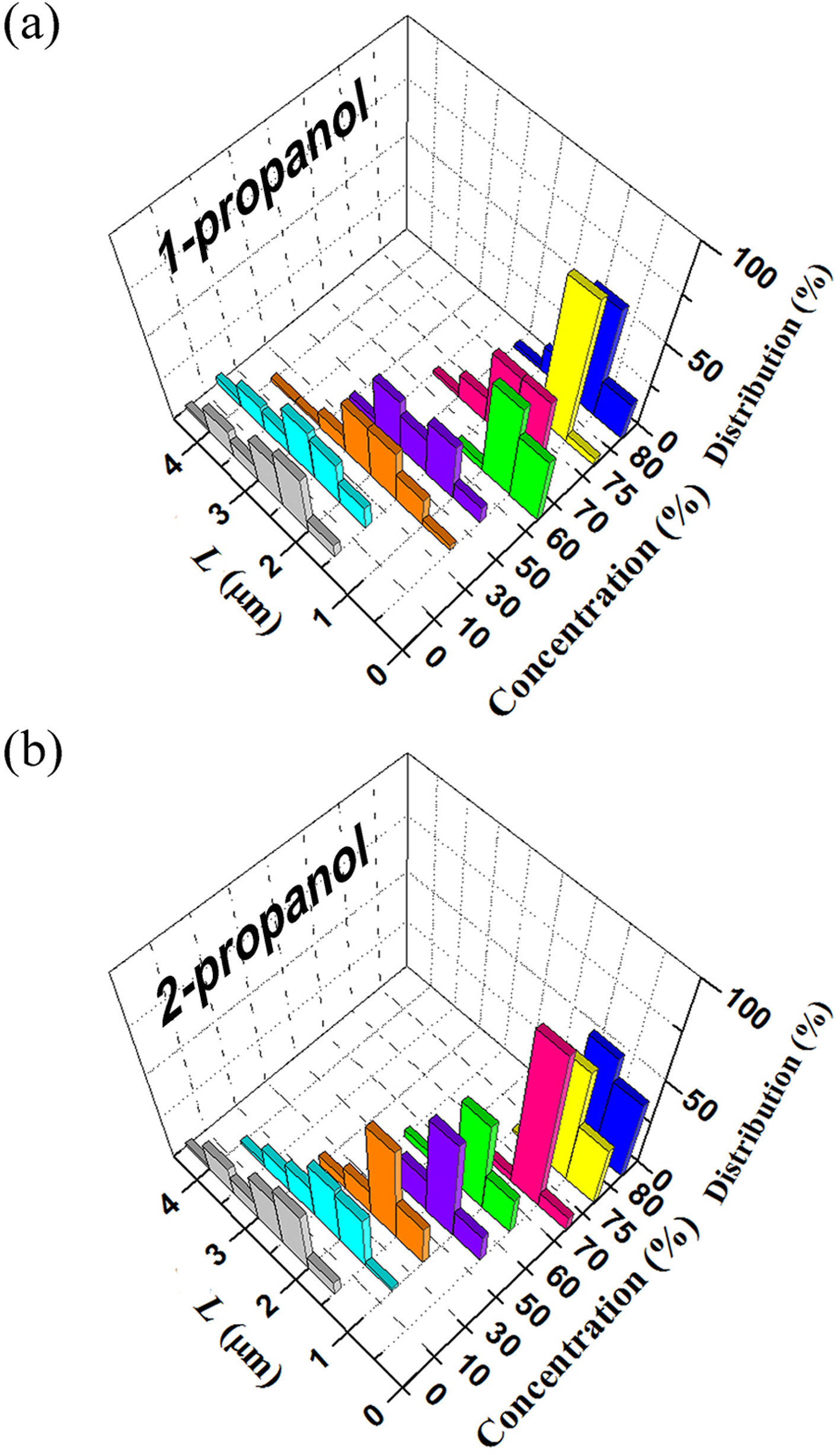
Histogram of the long-axis length *L* of λ-DNA molecules at different concentrations of (a) 1-propanol and (b) 2-propanol.

### Analysis of the Brownian motion of single DNA

As in Fig. 4, the trajectories of the center mass of individual single DNA indicate rather significant difference on the translational Brownian motion depending on the solution compositions: samples without propanol (Fig. 4 a), with 60% (v/v) (Fig. 4 b) and 70% (v/v) (Fig. 4 d) 1-propanol, and with 60% (v/v) (Fig. 4 c) and 70% (v/v) (Fig. 4 e) 2-propanol, and found large differences. The DNA molecule in 60% (v/v) 1-propanol solution is much more active than the other samples. The molecule without alcohol is less active than DNA in other solutions with alcohols. Due to the blurring effect of approximately 0.3 μm, the photos do not provide precise information on the actual sizes of individual DNA molecules, especially for the compact state. To gain insight into the effects of propanols on the size of DNA, we analyzed the Brownian motion of individual DNA molecules in different propanol solutions. From the motion trails of the center of mass of a single DNA molecule, we calculated the mean square displacement and evaluated the diffusion constant *D* for each DNA molecule using Eq.1: (36)

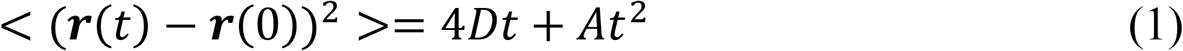

where ***r***(*t*) = (*r_x_, r_y_*) is the position of the center of mass for a DNA, < (***r***(*t*) − ***r***(0))^2^ > is the mean square displacement, and *A* is a numerical constant related to convective flow. The hydrodynamic radius *R_H_* is calculated from *D* based on the Stokes-Einstein relation, (37)

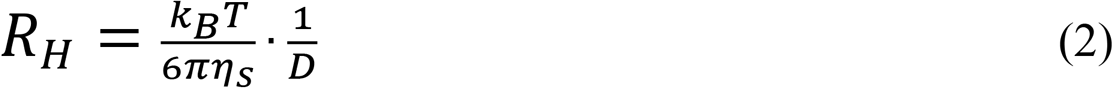

where *k_B_* is the Boltzmann constant and *η_s_* is the viscosity of the solvent at 298 K. The viscosity of the solvent was evaluated as described in the literature.(38)

**FIG. 4.**
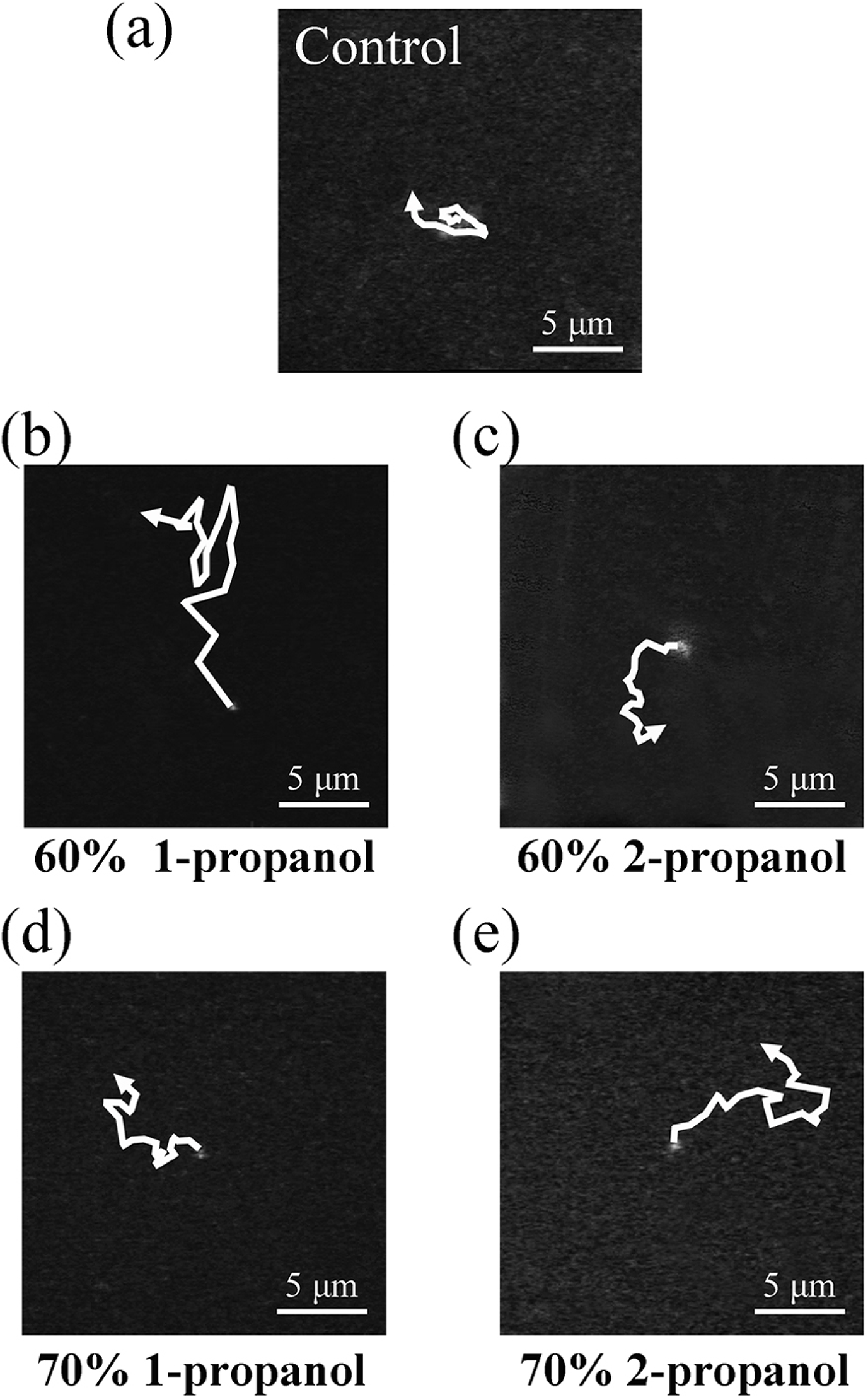
Trajectories of the center mass of individual molecules in (a) solution without propanol, (b) 60% (v/v) 1-propanol solution, (c) 60% (v/v) 2-propanol solution, (d) 70% (v/v) 1-propanol solution, and (e) 70% (v/v) 2-propanol solution observed by fluorescence microscopy for 3 seconds.

As shown in Fig. 5 c, the change in the hydrodynamic radius of DNA molecules, *R_H_*, exhibits similar trend as in the change in their long-axis length. For DNA molecules in different concentrations of 1-propanol solutions, *R_H_* decreased from 0.91 μm to 0.02 μm as the concentration of 1-propanol increased to 60 % (v/v). A minimum *R_H_* (0.02 μm) also appeared at 60 % (v/v). However, by changing the concentrations of 1-propanol from 60 %(v/v) to 75%(v/v), *R_H_* increases from 0.02 μm to 0.10 μm. This demonstrates that DNA molecules swell or decondense above 70 %(v/v). Whereas, in 2-propanol solutions, the *R_H_* of DNA decreased from 0.91 μm to 0.03 μm in a monotonous manner as the concentration of 2-propanol increased. The changes in the diffusion constant in Fig. 5 b thus clearly reveals the characteristic difference in the high-order structure of DNA with different propanol solutions in terms of the physico-chemical parameter of *R_H_*.

**FIG. 5.**
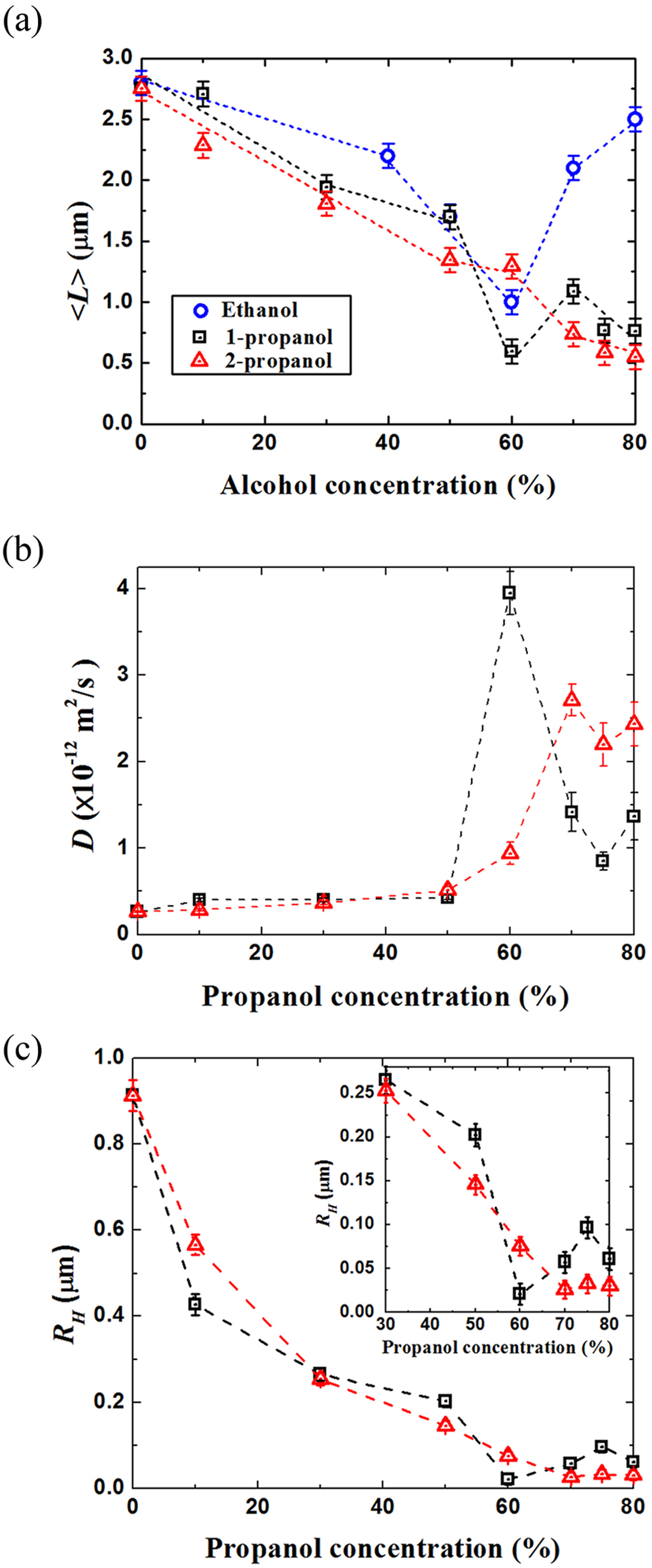
(a) The average long-axis length of DNA molecules at different alcohol solutions (19); (b) Diffusion constant of DNA molecules, *D*, at different propanol concentrations; (c) Hydrodynamic radius of DNA molecules, R_H_, in different propanol concentrations.

### Secondary structure of DNA molecules in alcohol solutions

As shown in Fig. 6 a, for DNA molecules in 1-propanol solutions, based on the positive band at around 275 nm and the negative band at around 245 nm, DNA maintained a B-like secondary structure(19) from 0% to 70% (v/v). The spectra approach zero when the concentration of 1-propanol is higher than 70% (v/v), which is attributed to the effect of precipitation accompanied by the condensation of DNA molecule. For samples in 2-propanol solutions (Fig. 6 b), the secondary structure changed to A-like form(19) from 30% to 60% (v/v), since the positive band is higher and the negative band is lower than those for samples in 0 % (v/v). The secondary structure returned to B-like at around 70% (v/v) of 2-propanol. At the concentration higher than 75% (v/v), the effect of DNA precipitation become non-negligible. To more clearly understand the observed change of the CD spectra, degrees of ellipticity (θ) at 270 nm are shown in Fig. 6 c. It is found that DNA molecules in 1-propanol only showed a B-like form before DNA deposition. On the other hand, in 2-propanol solution, both A-like and B-like forms appeared when the concentration of 2-propanol was lower than the DNA deposition concentration.

**FIG. 6.**
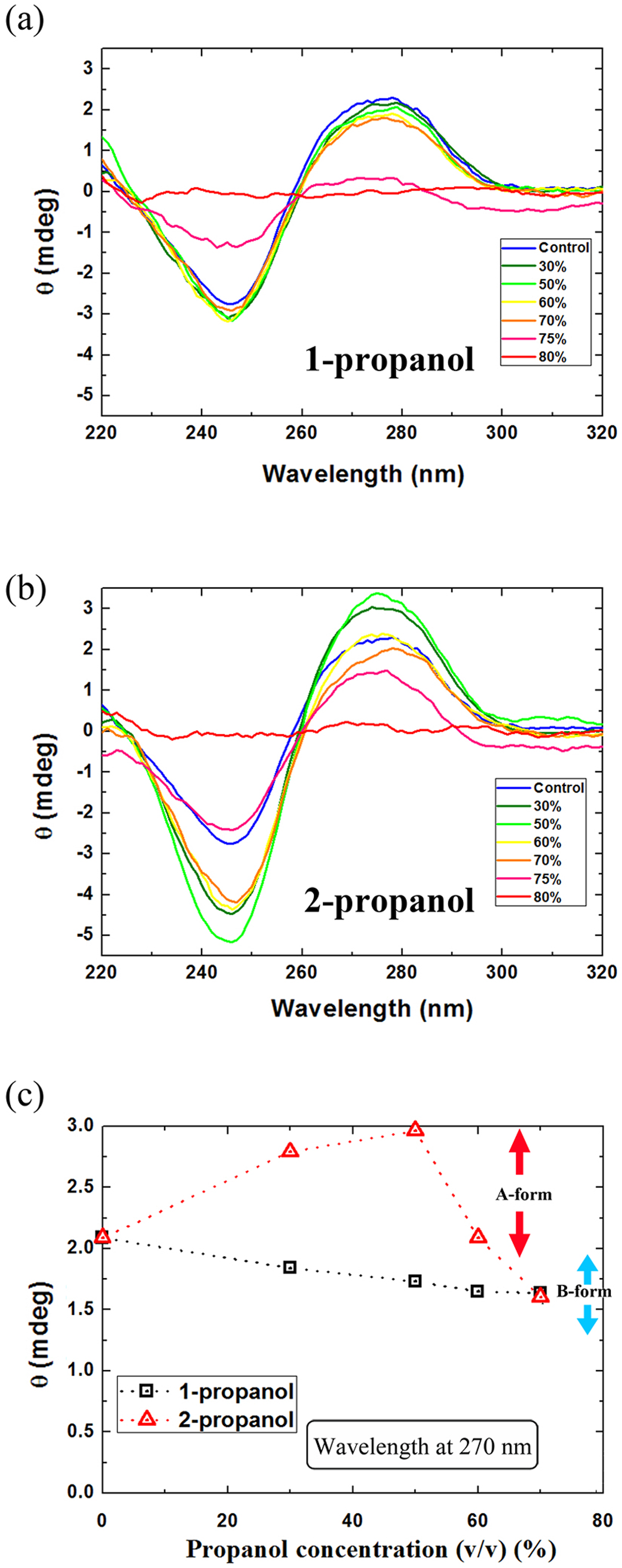
CD spectra of DNA (λ-DNA, 30 μM in nucleotide units) in (a) 1-propanol and (b) 2-propanol solutions; (c) Degree of ellipticity (θ) of CD spectra of DNA samples at 270 nm.

## DISCUSSION AND CONCLUDING REMARKS

From the experimental observation summarized in Fig. 5, it becomes clear that the long-axis length of DNA molecules in solution decreased as the concentration of 1-propanol increased to 60% (v/v), then increased slightly and remained constant as the concentration continued to increase. For 2-propanol solutions, the long-axis length decreased as the concentration increased, and then remained steady at a minimum value at high concentration. The hydrodynamic radius of DNA molecules calculated from Brownian motion showed essentially the same trends as the long-axis length of DNA as the concentrations of 1- and 2-propanol changed. As shown in Fig. 5 a, for DNA in solutions of ethanol, which is also commonly used in DNA precipitation, the long-axis length reached a minimum at 60% (v/v), then increased as the concentration continued to increase.(19)

DNA is negatively charged because of the negatively charged phosphate group in its structure.(39) Since 1-propanol molecules are straight-chain, like ethanol, they should exhibit similar polarity. The occurrence of the reentrant transition with ethanol and 1-propanol but not with 2-propanol may concern such geometrical difference of the chemical structure of the alcohols. The secondary structure of DNA molecules retained a B-like form in 1-propanol solutions. However, in 2-propanol solutions, it changed to an A-like form and then back to a B-like form as the concentration of 2-propanol increased. Moreover, past research by our group on the secondary structure of DNA in ethanol solutions demonstrated that DNA showed a B-, C- and A-like form as the concentration of ethanol increased.(19) The observed large difference in the effects of ethanol and propanol isomers on the DNA conformation will be discussed in relation to the nano-structure of an alcohol solution.(40–45)

The structural characterization of genomic DNA is one of the most important topics in medicine, biology and agricultural sciences. An indispensable procedure for the analysis of DNA in living cells is to isolate DNA molecules as precipitates from a crude mixture of a wide variety of cellular components. According to the standard experimental protocol in molecular biology and medicinal chemistry, the use of 2-propanol is recommended, but without any reasonable physico-chemical explanation for why 2-propanol is desirable.(4, 6) As a related phenomenon, we recently reported that ethanol causes a reentrant transition for DNA as its concentration increases. (19)

Past literatures have reported the formation of clusters in aqueous solutions of alcohols.(40–45) Dixit, et al.(40) performed a measurement of neutron-scattering on aqueous ethanol solution and reported the formation of nano-clusters rich in water molecules at high alcohol concentrations. The reentrant transition of the higher-order structure of DNA molecules is attributable to the association of water nano-clusters onto negatively charged phosphate groups along double-stranded DNA molecule. Through such hydration effect by water nano-clusters, the phosphate groups of DNA tend to dissociate to negatively charged state by eliminating counter cations to the nearby environment. Thus, DNA molecule undergoes conformational transition from compact state onto a swelled state at higher concentrations of 1 -propanol, similar to the reentrant transition of the higher-order structure of DNA depending on the concentration of ethanol.(19) As a next extension, it may of interesting to carry out the study to examine the different manner of mixing state between the propanol isomers, where water-rich nano-clusters will be generated with 1-propanol preferentially comparted to the solution with 2-propanol.

The long-axis length of DNA molecules, which is used to evaluate the higher-order structure, was significantly different between 1-propanol and 2-propanol solutions. It is found that, with an increase in the concentration of 1-propanol, DNA exhibits a reentrant transition, whereas 2-propanol causes a monotonous change in the DNA conformation. Based on this new insight into the difference in the physico-chemical effects of these propanol isomers, we discuss why 2-propanol solution is most frequently used(4, 6, 7, 9, 10) to retrieve genomic DNA molecules from living cells.

## AUTHOR CONTRIBUTIONS

Y.M., Y.Y. and K.Y. designed the research. Y.M. performed all experiments and analyzed the data. Y.M. and K.Y. wrote the article.

## ACKNOWLEDGEMENTS

This work was supported by JSPS KAKENHI Grant Numbers 15H02121 and 18J15422. Y. M. received support as a Research Fellow of the Japan Society for Promotion of Science.

